# Nucleosome repositioning in cardiac reprogramming

**DOI:** 10.1101/2024.11.05.622077

**Authors:** Sonalí Harris, Syeda S. Baksh, Xinghua Wang, Iqra Anwar, Richard E. Pratt, Victor J. Dzau, Conrad P. Hodgkinson

## Abstract

Early events in the reprogramming of fibroblasts to cardiac muscle cells are unclear. While various histone undergo modification and re-positioning, and these correlate with the activity of certain genes, it is unknown if these events are causal or happen in response to reprogramming. Histone modification and re-positioning would be expected to open up chromatin on lineage-specific genes and this can be ascertained by studying nucleosome architecture. We have recently developed a set of tools to identify significant changes in nucleosome architecture which we used to study skeletal muscle differentiation. In this report, we have applied these tools to understand nucleosome architectural changes during fibroblast to cardiac muscle reprogramming. We found that nucleosomes surrounding the transcription start sites of cardiac muscle genes induced during reprogramming were insensitive to reprogramming factors as well as to agents which enhance reprogramming efficacy. In contrast, significant changes in nucleosome architecture were observed distal to the transcription start site. These regions were associated with nucleosome build-up. In summary, investigations into nucleosome structure do not support the notion that fibroblasts to cardiac muscle cell reprogramming involves chromatin opening and suggests instead long-range effects such as breaking closed-loop inhibition.

## Introduction

In a previous report, we described the development of a set of tools to investigate nucleosome architecture (1). The approach differs from others in several ways. Firstly, the approach utilizes common software that is easily and readily available. In contrast, other approaches require the use of specific programs which pose significant barriers due to requirements to understand computer languages and a reliance on repositories which are not maintained. Secondly, and perhaps more importantly, our approach starts with specific groups of genes. The standard approach starts by investigating nucleosome architecture changes with the entire genome as a single group. Groups of genes are then identified that fit a pattern and biological information is subsequently extracted through ontological methods. A problem that we have found with gene ontology is there is often a mismatch between the constituent genes of gene ontological terms and the genes whose expression is relevant to the research question (1).

Once we developed our novel approach, we applied it to study nucleosome architectural changes during skeletal muscle cell differentiation. Three groups of muscle genes were chosen for study: those expressed solely in skeletal muscle; those expressed in both skeletal muscle and cardiac muscle; and those expressed solely in cardiac muscle. By way of a control, a fourth group was comprised of non-muscle genes. Each gene group was verified for specificity and was of sufficient size to avoid any potential bias from outliers. Our approach revealed that during myogenic differentiation, skeletal muscle and common muscle genes underwent specific and unique nucleosome changes not observed in the other groups. Nucleosomes in three regions (−2500bp to -2400bp, -650bp to -400bp and +600bp to +700bp) relative to the transcription start site shifted in a 3’ direction. In addition, there was notable a loss of nucleosomes surrounding the transcription start site.

Moving forward, we wanted to apply our approach to direct cardiac reprogramming. We, along with others, have reported that partial functionality can be restored to the infarcted heart by directly reprogramming scar tissue fibroblasts into heart muscle cells (2-5). The initial stages of direct reprogramming are characterized by several epigenetic events, notably loss of H3K27me3 and gain of H3K4me3 (6, 7). These epigenetic markers are believed to promote gene repression and gene activation, respectively, and are thought to work co-operatively to open chromatin, enabling RNApol-II to bind and initiate transcription (7). However, chromatin remodeling has not been proven. Moreover, other studies in different gene systems contend that epigenetic changes are merely bystander effects (8). The tools at our disposal enable us to address these questions further. As detailed in this report, direct reprogramming does induce changes to the nucleosome architecture specific to cardiac muscle genes. However, these changes occur at some distance from the TSS and appear to coincide with those observed in skeletal muscle genes during myogenic differentiation. Notably, there were no changes in the nucleosome architecture surrounding the TSS. Moreover, agents that are known to enhance reprogramming efficacy through enhanced gene expression also had no effect on TSS nucleosome architecture. In conclusion, our findings suggest that epigenetic events initiated by direct reprogramming do not open up chromatin for RNApol-II binding.

## Materials & Methods

### Databases

the RNA-seq(9) and MNase-seq(10) datasets are available on the NIH Single-Read Archive under accession numbers PRJNA1067687 and PRJNA837987 respectively.

### Gene groups

per our previous publication(1), genes were ascribed to specific gene groups.

The cardiac & common muscle gene group includes Actn2, Cacna1c, Kcna4, Kcnj2, Mb, Mef2C, Myh6, Myl2, Nebl, Ryr2, Scn5a, Sln, Tnni3, Tnnt2, and Ttn.

The skeletal muscle gene group includes Cacna1s, Myh1, Myh2, Myh3, Myh4, Myod1, Myog, Neb, Ryr1, Scn4a, Tnni1, and Tnni2.

The non-muscle group includes Atl1, Cd34, Cdh1, Col1a1, Col1a2, Dcx, Eno2, Eng, Flt1, Flt4, Map2, Mapt, Ncam1, Neurod1, Nlgn1, Pecam1, Postn, Rbfox3, Scn4a, S100a4, Syn1, Tek, Tcf21, Thy1, Vcam1, Vegfa, and Vwf. This group is larger to ensure sufficient coverage of alternative reprogramming trajectories (fibroblast, endothelial, neuronal).

### RNA-seq

analysis of our dataset (PRJNA1067687) was carried out as described. Briefly, neonatal cardiac fibroblasts were transfected with miR combo (reprogramming agent) or a control miRNA. Four days later, total RNA was extracted using the Quick-RNA MiniPrep Kit (Zymo Research, 11–328) and high-throughput sequencing was performed by the Duke Genomic Core on an Illumina HiSeq 4000. Gene expression was determined via bioinformatics programs within the Galaxy suite(11).

### Investigating nucleosome architecture

the tools used to investigate nucleosome architecture are described at length in our earlier report(1). The starting point was MNase-digested chromatin isolated from miR combo and control miRNA transfected neonatal cardiac fibroblasts and sequenced on a NovaSeq 6000 Kit (Illumina). The dataset has an accession number of PRJNA837987 in the NIH single-read archive(10). Bowtie was used to align sequences to the mouse genome and filtered for quality, true pairs and mitochondrial genome sequences. BamCoverage (1bp window, normalized to effective mouse genome size and in MNase mode) was used to determine read count for each promoter (−3kb upstream of the transcription start site to + 1kb downstream of the transcription start site) for the genes listed above. Data was exported as bedgraph and the bedgraph files were merged with the bedtools Merge BedGraph files tool. Once merged, read count data was normalized to ensure gene had equal weight via the equation:

For gene x, normalized read count_position-n_ = read count_position-n_/∑read count_genex_ where position-n is the position of the 1bp window with respect to the transcription start site.

To determine the effects on nucleosome architecture between two conditions the following formula was applied:

For gene x, Δnormalized read count_position-n_ = (read count_position-n_)myotube– (read count_position-n_)control.

Significances in Δnormalized read counts were determined by paired T-tests comparing normalized read counts in the two groups for each 1bp window.

### Statistics

Statistical methods are described in the figure legends.

## Results

Our approach for analyzing nucleosome architecture involves starting with a defined group of genes. A group of genes must be greater than 10 to ensure insensitivity to outliers (**Fig.1A**). Consequently, all gene groups were comprised of more than 10 genes.

**Figure 1.**
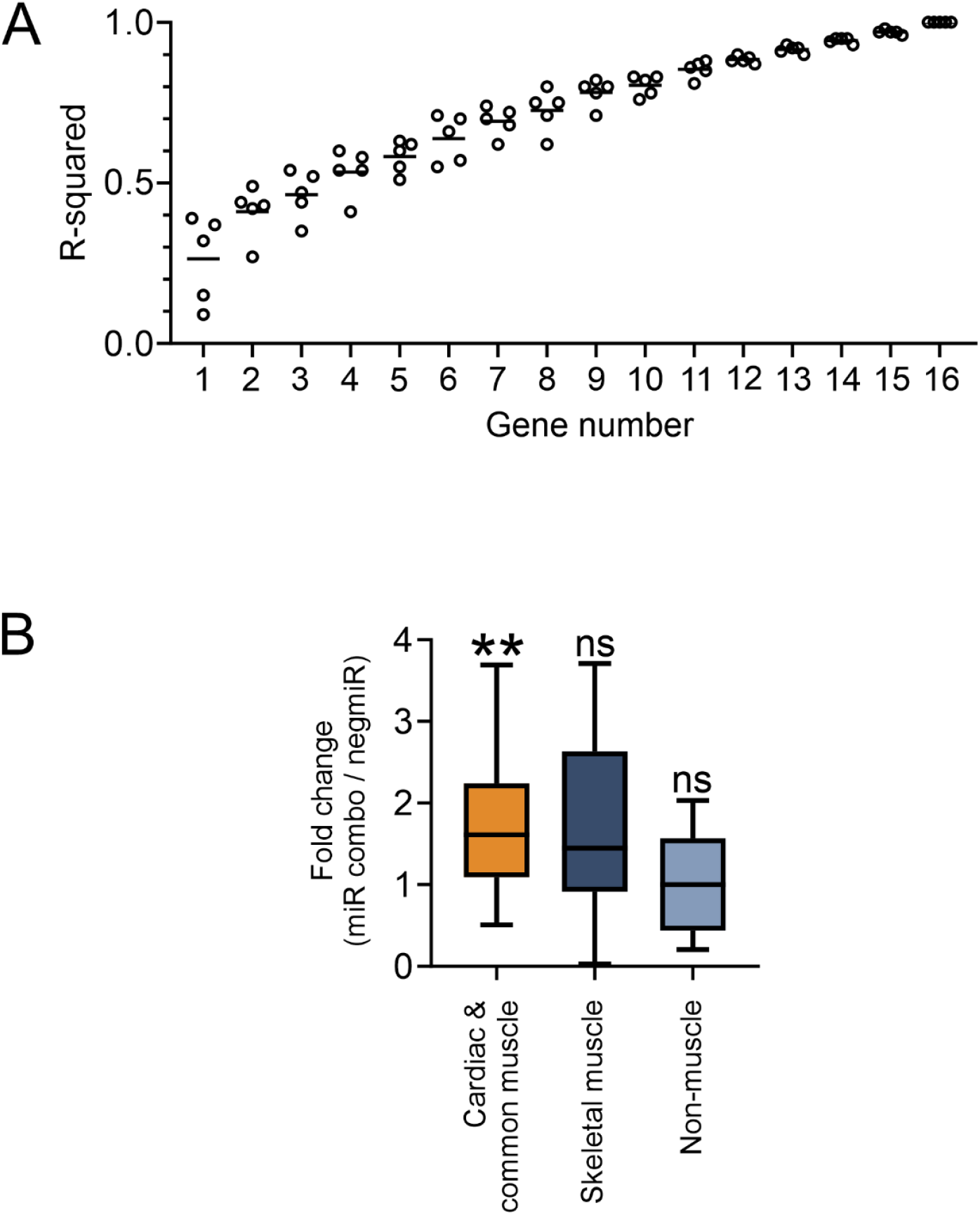
Expression of muscle and non-muscle genes during the direct reprogramming of fibroblasts into cardiac muscle cells. **(A)** To determine the minimum number of genes for analysis of nucleosome architecture, an experiment was conducted whereby genes were additively compared to their full-group. The comparisons were made on nucleosome density at a 1bp resolution across the full promoter and the R^2^ value noted after the addition of each gene. Five random sequences of genes were investigated. **(B)** RNA-seq data derived from fibroblasts four days after transfection with the reprogramming cocktail miR combo were investigated for the expression of the members of each listed gene-group. The data is expressed as a fold change in expression when compared to cells transfected with a control miRNA. N=3. Significances were determined by Wilcoxon Signed Rank Test (Pratt method, median = 1); ns – not significant, **P<0.01.

Our analysis of nucleosome architecture in fibroblasts to cardiac muscle cells focused on three groups of genes: (1) cardiac and common muscle genes; (2) genes specific to skeletal muscle; and, (3) non-muscle genes. In fibroblasts reprogramming to cardiac muscle cells, cardiac and common muscle genes are activated. In contrast, genes specific to skeletal muscle, as well as non-muscle genes, remain silent or are repressed (**Fig.1B**).

Regions of significant increase or decrease in nucleosome content were determined by comparing nucleosomes following transfection with the reprogramming agent miR combo and control miRNA. With respect to cardiac and common muscle genes, miR combo significantly altered nucleosome architecture in three regions of the promoter. These positions were -2600bp to -2300bp (Region A); -1800bp to -1400bp (Region B); and +350bp to +900bp (Region E) relative to the transcription start site (**Fig.2A**). In these regions there was a significant increase in nucleosome density. The efficacy of miR combo is enhanced by the addition of RNA-sensing receptor agonists (10, 12). The actions of these agonists on cardiac and common muscle genes was focused on a region -900bp to -550bp (Region C) relative to the transcription start site (**Fig.2A**). Neither miR combo alone, nor miR combo plus an RNA-sensing receptor agonist, had an effect on nucleosome content close to the transcription start site (−200bp to +200bp (Region D); **Fig.2A**).

**Figure 2.**
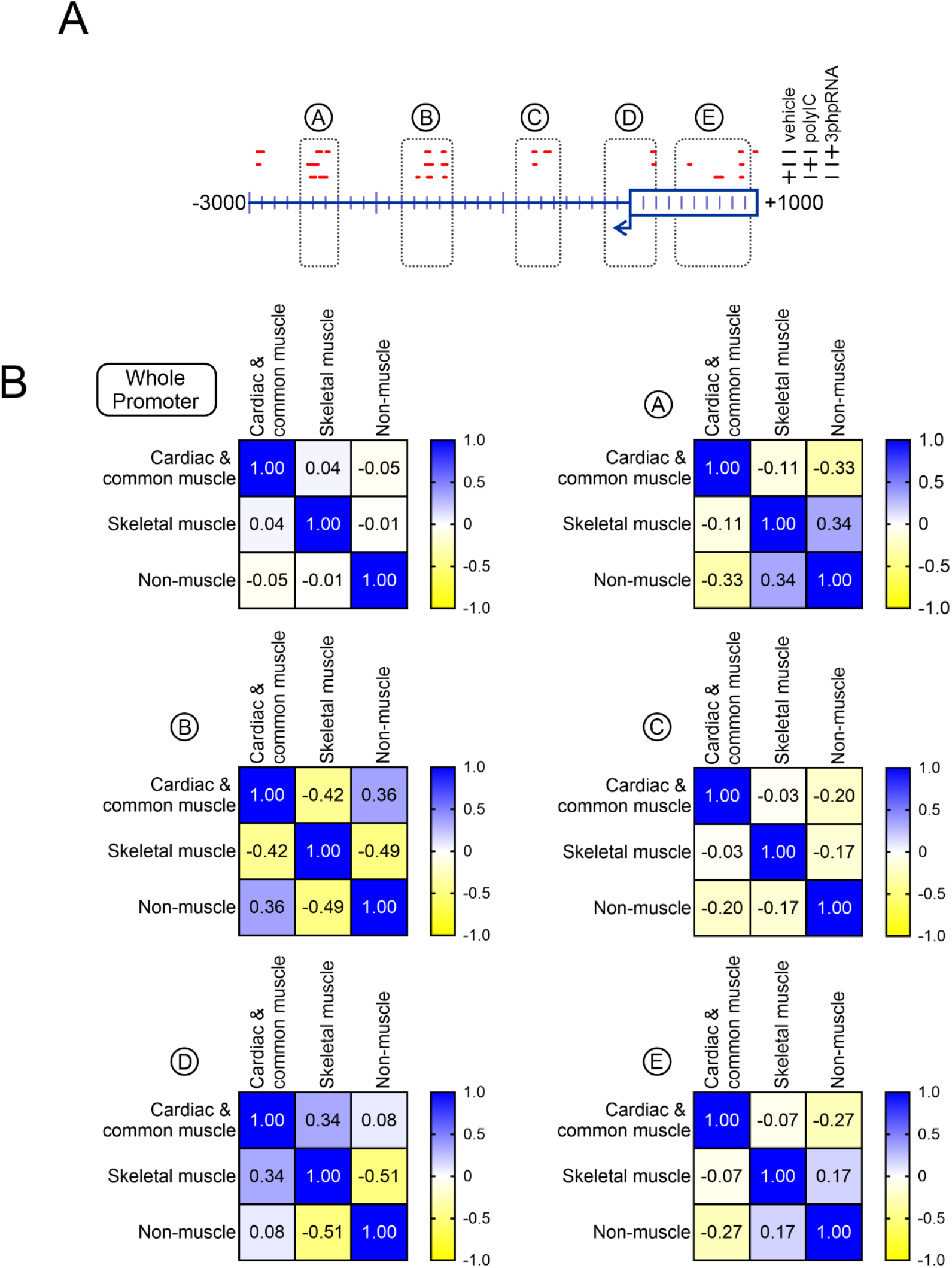
Direct reprogramming of fibroblasts into cardiac muscle cells is associated with significant changes in nucleosome architecture. **(A)** Our PRJNA837987 MNase-seq dataset was analyzed to determine the effects of direct reprogramming on nucleosome positioning. To compare nucleosome patterns within each group and across the three groups, the read counts were normalized by dividing the read count at each base-pair of the promoter by the sum of read counts across the promoter (a full description is provided in the methods). The schematic shows those regions in cardiac and common muscle gene promoters where nucleosomes were significantly enriched or depleted in response to reprogramming as well as in response to the reprogramming enhancers PolyIC and 3p-hpRNA. **(B)** Regions of significant enrichment or depletion (A, B, C and E) were analyzed further alongside a region encompassing the TSS (D) by determining the nucleosome correlations.

The regions described above were studied in more detail by comparing cardiac and common muscle genes with skeletal muscle and non-muscle genes. The comparisons were broken down into two measurements. Firstly, comparisons were made to determine if the test condition altered nucleosome position. Secondly, comparisons were made to determine if the test conduction induced nucleosomes to build up or are depleted. To address changes in nucleosome position, correlation matrices were determined. The R^2^-value represents how nucleosome positions change. For any given region, if nucleosomes in two genes occupy the same position relative to the transcription start site, they are in phase and the R^2^-value will be +1. Whereas, the R^2^-value will be -1 if the nucleosomes are fully out of phase (nucleosome in one group is completely absent in the comparator group). As shown in **Fig.2B**, in miR combo transfected cells, the position of nucleosomes across the whole promoter show no correlation between the three groups of genes. Correlations and anti-correlations are more evident in the aforementioned five regions of the promoter. Of the five regions, -2600bp to -2300bp (Region A), -900bp to -550bp (Region C) and +350bp to +900bp (Region E) are notable because they appear to define cardiac and common muscle genes from skeletal muscle and non-muscle genes on the basis of negative R^2^-values in the latter two gene groups (**Fig.2B**). Region -1800bp to -1400bp distinguishes cardiac and common muscle genes from skeletal muscle but not non-muscle genes and there is no difference in nucleosome positioning in cardiac and common muscle genes, skeletal muscle and non-muscle genes in the -200bp to +200bp (Region D) (**Fig.2B**).

In terms of nucleosome content, miR combo generally only affected cardiac and common muscle genes. While there was no effect over the whole promoter, miR combo induced significant nucleosome build-up in the -2600bp to -2300bp (Region A), -1800bp to -1400bp (Region B) and +350bp to +900bp (Region E) regions (**Fig.3**). The only significant event in the skeletal muscle and non-muscle groups was a significant increase in nucleosome content in the -900bp to -550bp (Region C) region of the latter group (**Fig.3**). Again, there was no significant change in nucleosome content for any group of regions in the region immediately adjacent to the transcription start site (Region D) (**Fig.3**).

**Figure 3.**
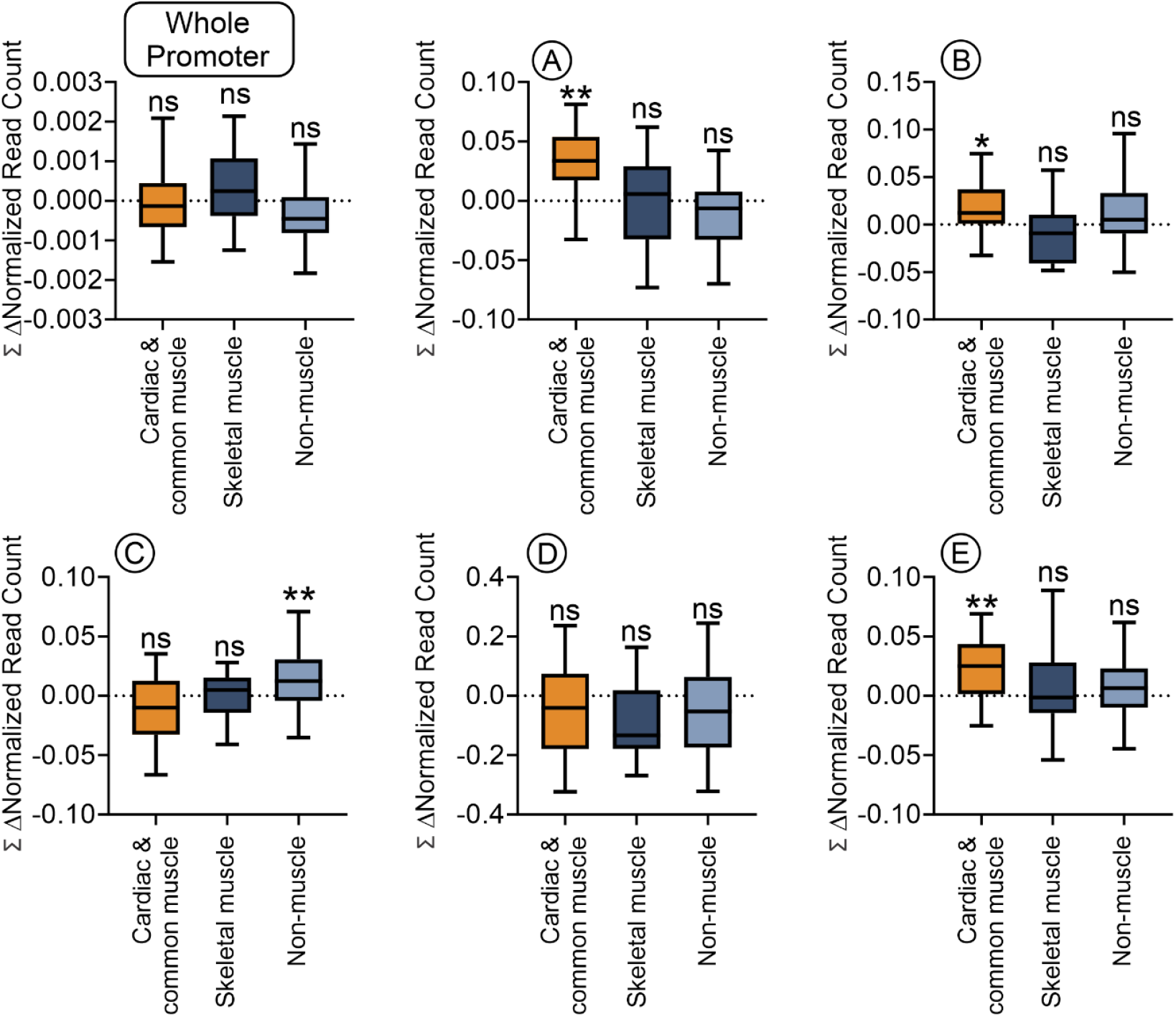
Cardiac and common muscle genes show significant nucleosome re-positioning away from the TSS. For each analyzed promoter, the change in read number (Δ normalized read count) following direct reprogramming (miR combo versus control miR) was calculated at a 1bp resolution and summed over the genomic region analyzed. Significances were determined by Wilcoxon Signed Rank Test (Pratt method, median = 0); ns – not significant, **P<0.01.

As mentioned above, fibroblast to cardiomyocyte reprogramming efficacy is improved by the addition of RNA-sensing receptor agonists(10, 12). RNA-sensing receptors are a broad group of proteins that are believed to share a common signaling pathway. We have found evidence for the role for two RNA-sensing receptors, TLR3 and Rig1. TLR3 and Rig1 appear to influence fibroblast to cardiomyocyte reprogramming through different pathways. We wanted to understand if TLR3 and Rig1 agonists potentially enhance reprogramming efficacy by changing nucleosome architecture.

Over the whole 4kb promoter, neither the TLR3 agonist PolyIC, nor the Rig1 agonist 3p-hpRNA influenced nucleosome content in any gene group (**Fig.4**). However, differences were noted in the analysis of the five aforementioned promoter regions. Again, the first comparisons were made to determine the effect on nucleosome movement by determining how in phase they were. In cardiac and common muscle genes, both PolyIC and 3p-hpRNA induced significant nucleosome re-arrangement in all five of the aforementioned promoter regions. Moreover, the effects of the two agonists differed from each other as evidenced by the low R^2^-values (**Fig.4**). The same trends were observed in skeletal muscle and non-muscle groups, whereby PolyIC and 3p-hpRNA induced nucleosome re-arrangement when compared to the control and the re-arrangements differed between the two ligands (**Fig.4**).

**Figure 4.**
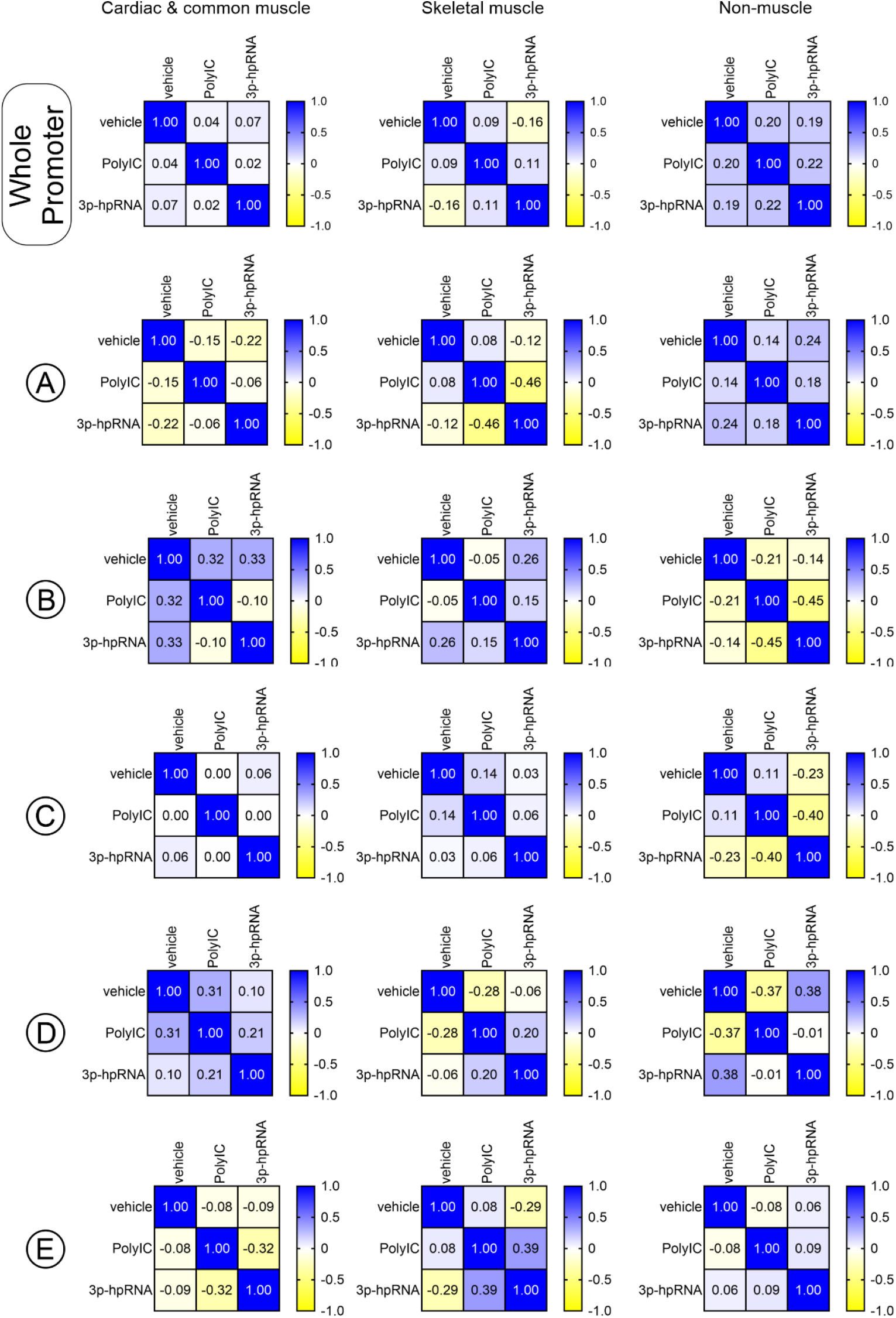
TLR3 and Rig1 agonists affect nucleosome architecture differently. For the whole promoter as well as regions A to E, the change in read number (Δ normalized read count) following direct reprogramming (miR combo plus PolyIC/3p-hpRNA versus miR combo) was calculated at a 1bp resolution and compared between the various gene groups. The matrices report the correlations between the various groups.

Following measurements of nucleosome movement, nucleosome content in the five promoter regions was quantified. Generally, both RNA-sensing receptor ligands induced the same trends. With respect to cardiac and common muscle genes, both PolyIC and 3p-hpRNA increased nucleosome content in the -1800bp to -1400bp and -200bp to +200bp regions while decreasing it in the -2600bp to -2300bp, -900bp to -550bp and +350bp to +900bp regions (**Fig.5**). Interestingly, PolyIC and 3p-hpRNA had the same effect on the equivalent regions of skeletal muscle genes (**Fig.5**). Many of the same features were also observed with non-muscle genes.

**Figure 5.**
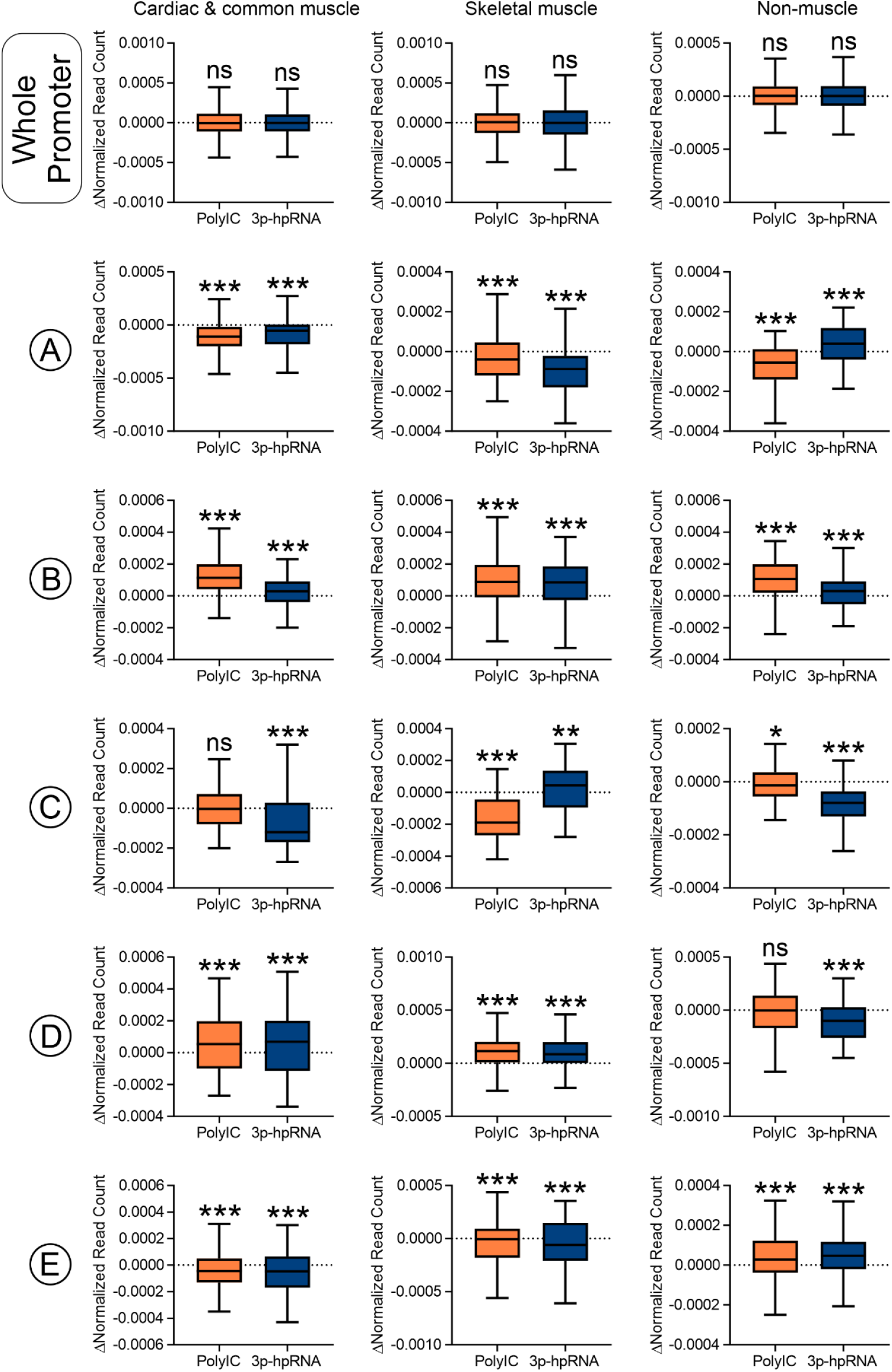
TLR3 and Rig1 agonists have opposing effects on nucleosome architecture in muscle and non-muscle genes. For the whole promoter as well as regions A to E, the change in read number (Δ normalized read count) following direct reprogramming (miR combo plus PolyIC/3p-hpRNA versus miR combo) was calculated at a 1bp resolution and summed. Significances were determined by Wilcoxon Signed Rank Test (Pratt method, median = 0); ns – not significant, *P<0.05, **P<0.01, ***P<0.001.

However, in contrast to muscle genes, PolyIC and 3p-hpRNA induced nucleosome loss in the -200bp to +200bp region and nucleosome gain in the +350bp to +900bp region (**Fig.5**).

## Discussion

When early events in the reprogramming of fibroblasts to cardiac muscle cells are studied’ there has been a focus on epigenetic changes. Indeed, early on there is a significant re-positioning of the epigenetic motifs H3K27me3, H2AK119ub and H3K4me3 (6, 7). H3K27me3 and H2AK119ub are typically labeled “repressive” while H3K4me3 is often labeled “activating”. They are believed to close and open chromatin respectively through the movement of nucleosomes. The idea is that open chromatin allows transcription factors to bind and activate specific genes. Due to the correlations between where these motifs are found and gene activity, researchers have assumed causality (7). However, this assumption is problematic because so-called repressive marks are also found on active genes and so-called activating marks are found on repressed genes (13-16). A further issue is one of universality. Antibody immunoprecipitations and sequencing are used to identify where the epigenetic motifs are located and biological function is subsequently inferred through ontology. While informative, gene ontological terms are curated lists and as such they can be incomplete, potential sources of bias, and not fully represent complex biological processes (17). What this means is that genes affected by reprogramming and do not fit the epigenetic paradigm have the potential to be missed. It was with these questions in mind that we developed an approach to study chromatin whereby we used novel analytical tools to identify significant changes in nucleosome structure. We first applied this approach to skeletal myoblast differentiation as this process is well characterized. Bearing in mind that gene ontological terms applied at the end of an analysis may miss important genes, we started with large panels of genes that are known to be specifically expressed in skeletal muscle cells. For controls, we also generated gene groups comprised of genes expressed solely in cardiac muscle and genes specific to alternate lineages. MNase-seq was used to measure nucleosome density at a single base-pair resolution over a 4kb region surrounding the transcription start site (−3kb to +1kb) of all the a priori chosen genes. After normalization to equally weight each gene and subtraction to calculate changes in nucleosome density, statistical methods were used to elucidate the effects of skeletal muscle differentiation on nucleosome structure (1). The approach we developed demonstrated that myogenic differentiation induced nucleosome loss immediately around the transcription start site of skeletal muscle genes. There was no such loss of nucleosomes around the transcription start site of cardiac muscle or non-muscle genes (1). These findings were in agreement with earlier studies (18) and demonstrated the validity our approach.

Having validated our approach in a model system, we have now applied it to study how nucleosomes respond to reprogramming. The expectation, based on the epigenetic data as well as the similarities between skeletal and cardiac muscle, was that nucleosomes would deplete from around the transcription start site of cardiac muscle genes. Unexpectedly this was not observed. Moreover, agents which enhance reprogramming efficacy by increasing cardiac muscle gene activity also had no effect on nucleosomes around the transcription start site. Instead, changes in nucleosome architecture were observed at some distance from the transcription start site. There were three significant regions specific to cardiac muscle genes: - 2600bp to -2300bp (Region A), -900bp to -550bp (Region C) and +350bp to +900bp (Region E) relative to the transcription start site. Rather than showing nucleosome depletion, as might be expected, all three regions showed significant nucleosome accumulation. Taken together, this data is hard to square with the aforementioned model of repressive and activating epigenetic motifs regulating chromatin opening to modify gene expression. Analyses of nucleosome architecture in fibroblast to cardiac muscle reprogramming are hard to come by and we are aware of only one other report where the authors studied nucleosome architecture via the alternative ATAC-seq method (19). Like our study, they found that the majority of changes in nucleosome occupancy were at some distance away from the transcription start site (19). Moreover, the authors provided evidence that changes in nucleosome occupancy were due to transcription factors binding to chromatin. Long distance regulation of gene activity, as suggested in both studies, could occur through mechanisms such as breaking closed-loop inhibition, whereby nucleosome architecture is such that the gene is silenced through loop formation (20). The further implication from both studies is that epigenetic marker re-positioning are not causative agents in reprogramming, but are more to stabilize the gene activity patterns (8).

We have reported that the efficacy of reprogramming fibroblasts to cardiac muscle cells is increased by RNA-sensing receptor agonists. To date, we have found evidence for the involvement of two RNA-sensing receptors: TLR3 (12) and Rig1 (10). Both TLR3 and Rig1 enhance efficacy by increasing cardiac muscle gene activity. However, the two receptors achieve this effect differently. While TLR3 utilizes the transcription factor NFkB, Rig1 instead utilizes the transcription factor YY1. We were interested if this difference in transcription factor utilization was mirrored in differences in nucleosome structure. Overall, both TLR3 and Rig1 agonists had a significant impact on nucleosome architecture. Interestingly, and mirroring their disparate transcription factor utilization, TLR3 and Rig1 agonists induced the adoption of distinct nucleosome architectures as there was little correlation in terms of nucleosome positions. TLR3 and Rig1 agonists appeared to affect muscle and non-muscle genes differently. While both agonists induced nucleosome depletion on muscle genes, the opposite was found on non-muscle genes. Thus, TLR3 and Rig1 agonists appear to enhance muscle gene activity through nucleosome re-arrangement and depletion.

In summary, studies of nucleosome architecture suggest that in the context of cellular reprogramming cardiac muscle genes are activated via changes in nucleosome positioning distal to the transcription factor start site.

## Acknowledgements

The work was supported by the Edna and Fred L. Mandel, Jr. Foundation as well as the National Heart, Lung and Blood Institute (R01 HL131814-01A1).

